# Social Context and Drug Cues Modulate Inhibitory Control in Cocaine Addiction: involvement of the STN evidenced through Functional Magnetic Resonance Imaging

**DOI:** 10.1101/2023.09.06.556336

**Authors:** Damiano Terenzi, Nicolas Simon, Michael Joe Munyua Gachomba, Jeanne Laure de Peretti, Bruno Nazarian, Julien Sein, Jean-Luc Anton, Didier Grandjean, Christelle Baunez, Thierry Chaminade

## Abstract

**Background:** Addictions often develop in a social context, although the influence of social factors did not receive much attention in the neuroscience of addiction. Recent animal studies suggest that peer presence can reduce cocaine intake, an influence potentially mediated, among others, by the subthalamic nucleus (STN). However, there is to date no such neurobiological study in humans.

**Methods:** This study investigated the impact of social context and drug cues on brain correlates of inhibitory control in individuals with and without cocaine use disorder (CUD) using functional Magnetic Resonance Imaging (fMRI). Seventeen CUD participants and 17 healthy controls (HC) performed a novel fMRI Stop-Signal task in the presence and absence of an observer while being exposed to cocaine-related and neutral pictures as cues.

**Results:** The results showed that CUD participants, while slower at stopping with neutral cues, recovered control level stopping abilities with cocaine cues, while HC did not show any difference. Neuroimaging revealed the involvement of frontal cortical regions and of the subthalamic nucleus (STN) in inhibitory control, which was modulated by both social context and drug cues in CUD participants but not in HC.

**Conclusions:** These findings highlight the impact of social context and drug cues on inhibitory control in CUD and suggest potential targets for intervention such as the STN, also emphasizing the importance of considering the social context in addiction research and treatment.

## 1. Introduction

Cocaine addiction, like most other addictions, takes place in a social environment that may influence drug use (1). Although this influence has long been suggested by social science studies (2–4), it is only recently that neurobiological studies have started to consider the role of social factors such as the others’ presence, either as an alternative reward to substances of abuse (5–7) or as a constant part of the consumption social environment (8,9).

Importantly, social factors may have opposite effects on drug-taking depending on the drug used such as psychostimulants (cocaine) or depressant (alcohol), but also depending on the experimental conditions. It seems indeed that in animals, the presence of peers can decrease the self-administration of morphine (10) or cocaine (9), whereas it can increase the self-administration of nicotine (11), ethanol (12), amphetamines (13), or cocaine in different conditions (8). Interestingly, in a recent study in rats, the optimal condition to reduce cocaine consumption, compared to other peers (familiar and/or exposed to cocaine), was the presence of an unfamiliar peer naive to cocaine. Similarly, in humans with psychostimulant use disorder, similar beneficial effects of the peer presence were observed in a cross-sectional survey assessing the social environment at the moment of stimulant use (9).

Lesioning the subthalamic nucleus (STN) in rats, a small lens-shaped nucleus that belongs to the basal ganglia, potentiated the beneficial effect of peer physical presence or playback of ultrasonic vocalizations of a stranger peer, thus positioning the STN as a key neurobiological substrate of social influence on drug intake (9,14,15). Furthermore, there are evidence to suggest STN deep brain stimulation (DBS) as a therapeutic strategy for addiction (16–20) which even led to one clinical attempt to date (21). There is currently no study that has combined social context manipulations in human addiction with the study of its neural correlates.

In human studies, the Stop-Signal task (SST) is one of the most used measures of impulsivity of action (22) as it requires inhibition of an overlearned ongoing action. Neuroimaging studies employing the SST have shown that the activity in the STN increases when participants successfully inhibit their response to the stop signal (23– 25). In contrast, lesioning the STN in rats leads to impaired ability to stop an ongoing action (26) and affects other forms of impulsivity of action (27). The STN receives direct inputs from many cortical areas (the motor cortex, the inferior frontal gyrus (IFG), the orbitofrontal cortex (OFC), Anterior Cingulate cortex (ACC), constituting the so-called hyperdirect pathways. These cortico-STN hyperdirect pathways facilitate the suppression of the ongoing motor response (28–30). Studies employing the SST in individuals with Parkinson’s disease (PD) who received STN DBS have further provided causal evidence for the involvement of the STN in successful inhibitory control (31–33).

The SST has found application in addiction research, as the ability to resist drug seeking and consumption rely heavily on voluntary, deliberate inhibition (34). Previous neuroimaging studies have revealed an aberrant response inhibition in addiction, which is accompanied by altered functional activity in frontal regions including the inferior frontal gyrus (IFG), the orbitofrontal cortex (OFC), and the dorsolateral prefrontal cortex (dlPFC) (35–41). A recent fMRI study on cocaine addiction has delved deeper into investigating how drug cue salience (in the form of words) affects the modulation of inhibitory control while participants performed the SST. The findings revealed that these cues have an impact on the neural patterns associated with inhibitory control in PFC regions (42).

Since addictive behavior can be considered as the result of a lack of inhibitory control leading to compulsive drug use (34,42,43), in this study we employed a SST to specifically engage brain regions involved in inhibitory control, such as the STN and frontal cortical areas directly connected to it: the IFG and OFC. These brain regions are also involved in reward processing(44–46). Hence, we expect that these brain areas may be strongly involved in our task and in our experimental conditions. In particular, we aimed to examine whether the social context could influence inhibitory control in individuals with cocaine addiction. To do so, we developed a novel fMRI version of the SST, namely the “social SST”. Participants with cocaine use disorder (CUD) and healthy controls (HC) performed the experimental task under two social contexts: in half of the 4 sessions, they were made to believe that they were observed by another person they could see on a video screen in the scanner (observing condition) and in the other half, no observer was present (non-observer condition). Furthermore, to induce some form of cocaine craving, we used cocaine-related stimuli (compared to neutral control stimuli) as cues indicating beginning and end of individual trials. These cues were chosen in order to trigger arousal, anticipation, and changes in behavioral motivation (43,47–50), as well as to affect inhibitory control in drug addiction (42).

We expected that inhibitory control in CUD would be affected by both drug cues stimuli and social context and that this may be mediated by brain regions including the STN and frontal areas such as OFC and IFG.

## 2. Materials and methods

### 2.1 Participants

The study included right-handed individuals aged 18 to 65. Seventeen participants with cocaine use disorder (CUD participants; 4 females /13 males, mean age 35.5, SD 9.23) and seventeen matched healthy controls (HC participants; 9 females/ 8 males, mean age 31.8, SD 13.0) took part in the study. CUD participants were recruited from the addiction unit of the Timone University Hospital in Marseille (France). All met DSM-V criteria for current cocaine addiction and underwent the Mini International Neuropsychiatric Interview (MINI) (51). Craving and withdrawal symptoms of CUD participants were determined using the Cocaine Craving Questionnaire (CCQ) (52). See Table 1. One of the CUD participants did not complete the CCQ.

Exclusion criteria consisted of i) a history of major psychiatric or neurological disorders, ii) MRI contraindications, iii) the use of psychotropic drugs, iv) and the absence of cocaine drug in urine assay (for CUD participants). The two groups of participants were matched for demographic variables such as age (*U* = 192.5, *p* = 0.101) and gender (χ^2^(1) = 3.11, *p* = 0.08) but difference for their levels of education reached significance (*U* = 72.5, *p* = 0.012). The study was in accordance with the Declaration of Helsinki and approved by the local Ethics Committee and the national CPP (under the license #2017T2-34). All participants gave written informed consent to participate in the study.

**Table 1.** Demographic, clinical and questionnaire data mean (and standard deviations) in the Cocaine Use Disorder and Health Control participants. CCQ = Cocaine Craving Questionnaire; DSM-V = The Diagnostic and Statistical Manual of Mental Disorders, Fifth Edition; SUD = Substance use disorder.

|  | CUD<br>(n=17) | HC<br>(n=17) |
| --- | --- | --- |
| Gender (female) | 4 | 9 |
| Age (y) | 35.5 (9.2) | 31.8 (13.0) |
| Education (y) | 14.5 (2.4) | 16.6 (2.1) |
| Duration of cocaine use (y) | 13.7 (7.5) | - |
| CCQ | 28.8 (13.9) | - |
| DSM-V SUD | 7.3 (2.3) | - |
| Cigarette use per day (cigarettes) | 7.6 (7.0) | - |

### 2.2 Social Stop Signal Task

In a standard SST, participants are instructed to respond as quickly as possible to a common neutral Go signal but must inhibit their response when this signal is followed by a rare Stop-signal (22.5% of trials), typically presented as an auditory tone or visual cues superimposed onto the Go signal (23,53). In our modified version of the SST, we utilized cocaine-related and neutral control pictures as cues to simulate real situations involving cocaine craving for CUD subjects. These cues have been shown to elicit arousal, anticipation, and changes in behavioral motivation (43,47,54) and can impact inhibitory control in individuals with drug addiction (42).

During the task performed in the scanner, participants were required to press a button using their index or middle finger of the right hand as quickly as possible upon the presentation of a left or right arrow (Go trials). The arrow appeared superimposed on the cue, which could be either neutral or cocaine-related, allowing us to investigate the potential impact of cocaine-related cues on inhibitory control. During Stop trials, following the arrow presentation, a stop signal (sound) occurred after a variable delay (stop-signal delay; SSD). Participants were instructed to withhold their response upon hearing the stop signal (Figure 1B). The SSD duration was set at 250 milliseconds and then adjusted independently for cocaine and neutral stimuli based on the participant’s success at stopping. Successful stops led to a 50 millisecond increase in the SSD duration for subsequent trials, increasing the difficulty. Conversely, unsuccessful stops resulted in a 50-millisecond decrease in the SSD duration, making subsequent trial easier (23).

Participants completed four runs of the task, with each run consisting of 12 blocks of 10 trials. Each run lasted approximately 5 minutes, for a total of 20 minutes. Each block was indicated with GO or STOP written instruction, indicating whether or not there could be stop signals occurring approximately in 20% of trials within the block. There was a total of 480 trials, with 108 Stop trials (22.5% of the trials). Importantly, to examine the impact of the presence of an observing person on inhibitory control in cocaine addiction, participants were led to believe that they could be observed via a camera by the experimenter TC wearing a labcoat in the adjacent scanner control room. In two runs they saw a visual feedback of the observing person displayed onscreen (observer condition), while the 2 other runs presented the video with an empty chair (non-observer condition). However, in reality, both videos shown to them were prerecorded to ensure rigorous control of the social context across participants. The first run always included the observer, to reinforce the cover story that explained to them that “the experimenter will observe you, especially at the beginning of the experiment, to ensure you understood and perform the task correctly”, and the order of the remaining 3 runs was randomized.

The three main factors of the experimental paradigm are the experimental group (CUD vs. HC), an inter-subject factor, while the cue type (cocaine vs. neutral) and social context (observer vs. no-observer) are intra-subject factors.

**Figure 1.**
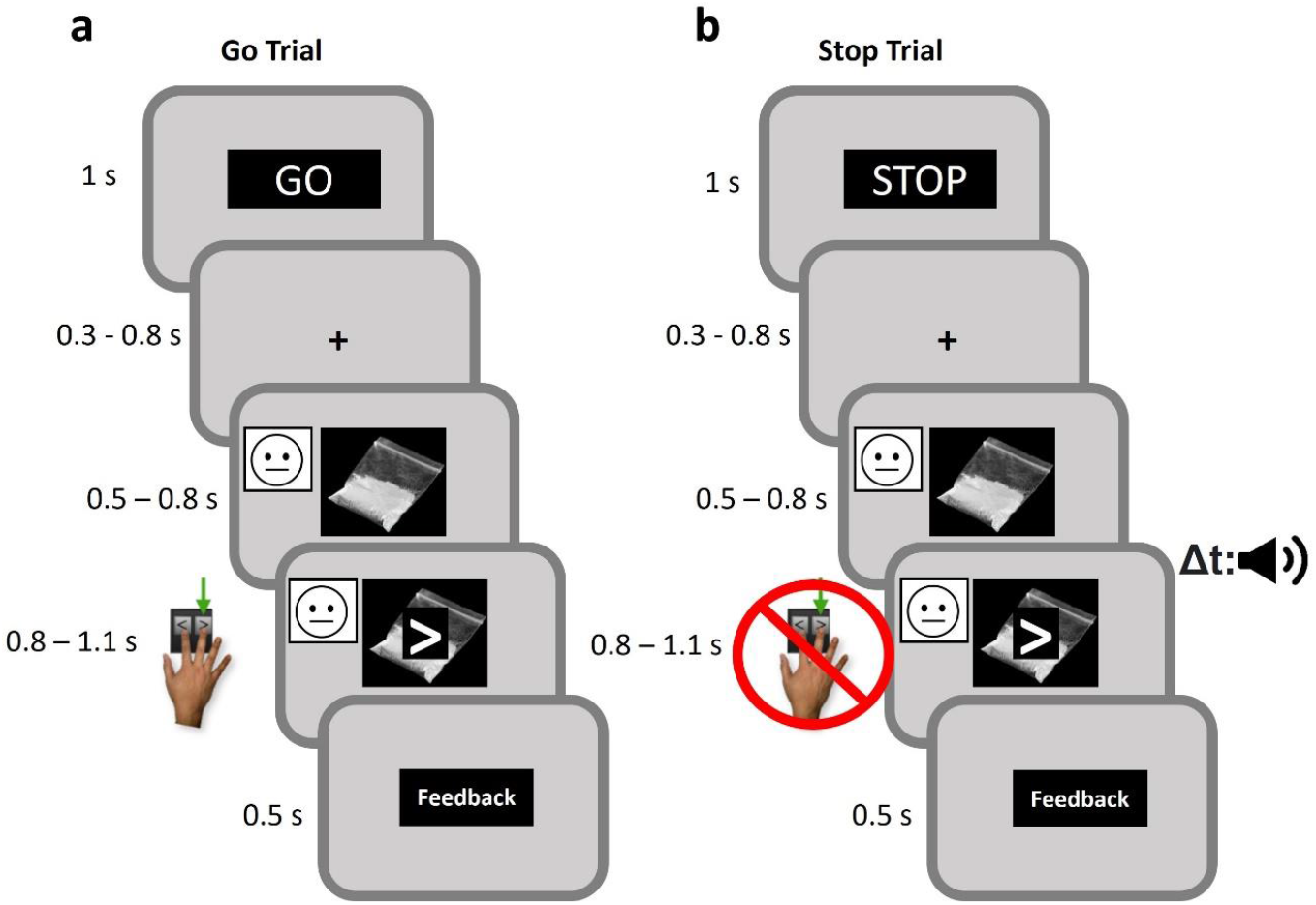
Schematic representation of the Stop Signal Task. Example of the observer condition. Each trial starts with a fixation point (+) (jittered duration between 0.3 and 0-8 s) followed by a neutral or cocaine-related cue (jittered duration between 0.5 and 0-8 s). In a Go Trial (A) the participant must press a button with the index or the middle finger of the right hand as quickly as possible after the presentation of an arrow pointing left or right, respectively (the maximum RT is 1.2 s). The arrow is superimposed on a cue which can be either neutral or cocaine related. Conversely, in a Stop trial (B), after the arrow and cue presentation, a sound (stop signal) occurs after a variable stop signal delay (SSD) indicating the participants to withhold their response. Each trial ends with a feedback screen indicating the accuracy of the participant’s response.

### 2.3 Data analyses

#### Behavioral data

Data were analyzed using R software version 4.1.2 (https://www.r-project.org/). Accuracy (the probability of giving a correct response) in Stop and Go trials was predicted with two separate logistic mixed effects models using the lmer function in the lm4 package (55), and explored using the Anova function type 3. Predictors consisted of the three experimental factors, group (CUD / HC), cue (cocaine / neutral), and the social context (observer / no observer). Participants’ ID was used as a random intercept.

We further predicted Reaction Times (RT) in correct Go trials, as well as Stop Signal Reactions Times (SSRT) in correctly inhibited Stop Trials using linear mixed models (LMMs). Predictors were the same as the ones described above. SSRT, which is the time required for one to successfully stop an ongoing response (i.e. not pressing the button) cannot be directly measured and was thus estimated using the Race model (23,56). More in details, RT on Go trials were rank ordered. Then, the *n*th RT was selected, where *n* is the result of the multiplication of the number of Go RTs by the probability of giving a response at a specific SSD. To get the SSRT, SSD was subtracted from this value. This process was repeated for each one of the central SSDs (35) and for each participant in each experimental condition (observer vs. non observer; cocaine-cue vs. neutral-cue). When predicting SSRT, the SSD was used as covariate in the model to account for SSD related differences in the SSRT.

#### Neuroimaging data

Data were acquired with a 3T MRI system (Siemens Magnetom Skyra) using a 64-channel head coil. Standard procedures were employed to preprocess the fMRI data. The volumes acquired correspond to the blood oxygenation level-dependent signal (BOLD) in 2.5 × 2.5 × 2.5 cm^3^ voxels of the brain (repetition time 1.224 s). Each volume includes 54 slices of 84 × 84 in-plane voxels. The volumes were slice-time corrected, realigned on the first one, and corrected for the deformation due to the local distortion of the magnetic field and participants’ movement. Spatial normalization of the imaged brains of all participants in the standard MNI space was performed using the DARTEL procedure (57). Individual analyses were performed for each participant and run.

The following events were modeled after convolution with a canonical hemodynamic response function: Stop Correct Cocaine, Stop Correct Neutral, Stop Incorrect Cocaine, Stop Incorrect Neutral. Eventually, a final event grouped all trials that did not fit in the previous categories, such as failed go trials (e.g. answer given on the wrong side or absence of recorded click). Trials were modeled as events, with null duration and onset at the presentation of the arrow. Several nuisance covariates were calculated to delete motion and physiological artifacts using the RETROICOR method from the heart pulse and respiration monitored using a photoplethysmograph and pneumatic belt, respectively, global grey matter signal, white matter activity, and cerebrospinal fluid activity (PhysIO toolbox from the TAPAS toolkit) (58). Single regressors represented the volumes with large movements from the participant.

To examine the neural signature of inhibitory control during the SSRT (Aron and Poldrack, 2006), contrasts between Stop Correct and Stop Incorrect (42,59) were computed for each Cue type (Cocaine / Neutral) and used in a second level analysis performed with GLM-Flex. GLM-Flex toolbox (https://habs.mgh.harvard.edu/researchers/data-tools/glm-flex-fast2/) allowed the analysis of factorial designs when more than 2 factors are present, and the effects of between-participants variable Group, between-run factor Social Context and within-run factor Cue Type were modelled. The three-way interaction Group x Social Context x Cue Type was computed, while factors Participants and Sessions were used as between- and within-participant random factors respectively.

## 3. Results

### 3.1 Behavioral results

#### Stop Signal Reaction Times (SSRT) in Stop trials

We first checked the assumption of the Race model (Verbruggen et al., 2019) according to which mean RT in incorrect Stop trials should be faster than mean RT in correct Go trials. Our results revealed indeed this difference (t(33) = 2.65, p = 0.012). Next, we performed a LMM investigating SSRT in correctly inhibited Stop trials. We limited our initial analyses to trials with neutral cues to maintain consistency with prior stop-signal task studies not using cocaine cues. This analysis showed a main effect of Group [χ^2^ (1) = 86.38, p = 0.028]. Post-hoc analysis showed slower SSRT (reduced inhibition) for users compared to controls (β = -37.26, t = -2.05, *p* = 0.048). No other significant effects of factors emerged (all *Ps* > 0.439). When performing a second LMM including all trial types (both neutral and cocaine cues), the two groups were no longer different in their SSRT. In addition, this model led to a significant two-way interaction Group x Cue type [X^2^ (1) = 6.38, *p* = 0.011]. Post-hoc analysis showed that cocaine users had faster SSRT for cocaine-related trials compared to neutral ones (β = -22.24, t = -2.71, *p* = 0.035) (Figure 2). This result persisted even when controlling for participants’ educational level (β = -22.23, t = -2.71, *p* = 0.035).

#### Accuracy in Stop trials

The mixed effects logistic regression model investigating the accuracy on Stop trials revealed no significant main effects of Group [χ^2^ (1) = 1.72, p = 0.189], Cue type [χ^2^ (1) = 0.46, p = 0.496], Social context [χ^2^ (1) = 0.49, p = 0.485]), nor significant interactions between these factors (all *Ps* > 0.250).

#### Accuracy in Go trials

the mixed effects logistic regression model investigating the accuracy on Go trials showed no significant main effects of Group [χ^2^ (1) = 1.68, p = 0.194], Cue type [χ^2^ (1) = 1.26, p = 0.261], Social context [χ^2^ (1) = 0.84, p = 0.358]; while the triple interaction Group x Social context x Cue type resulted statistically significant [χ^2^ (1) = 4.71, p = 0.030]. Post-hoc analysis (Tukey correction) showed no significant results (all *Ps* > 0.075).

#### Reaction Times (RT) in Go trials

The LMM for RT in correct Go trials showed a marginally significant main effect of Group [χ^2^ (1) = 3.82, p = 0.0506] and a significant two-way Group x Cue type interaction [χ^2^ (1) = 4.46, p = 0.034]. There were no other significant results (all *Ps* > 0.20). Post-hoc analysis on the main effect of Group showed that cocaine users were slower than control participants in Go trials (β = 45.3, t = 2.20, *p* = 0.028). Post-hoc analysis on the Group x Cue type interaction did not reveal significant results (all *Ps* > 0.078).

**Figure 2.**
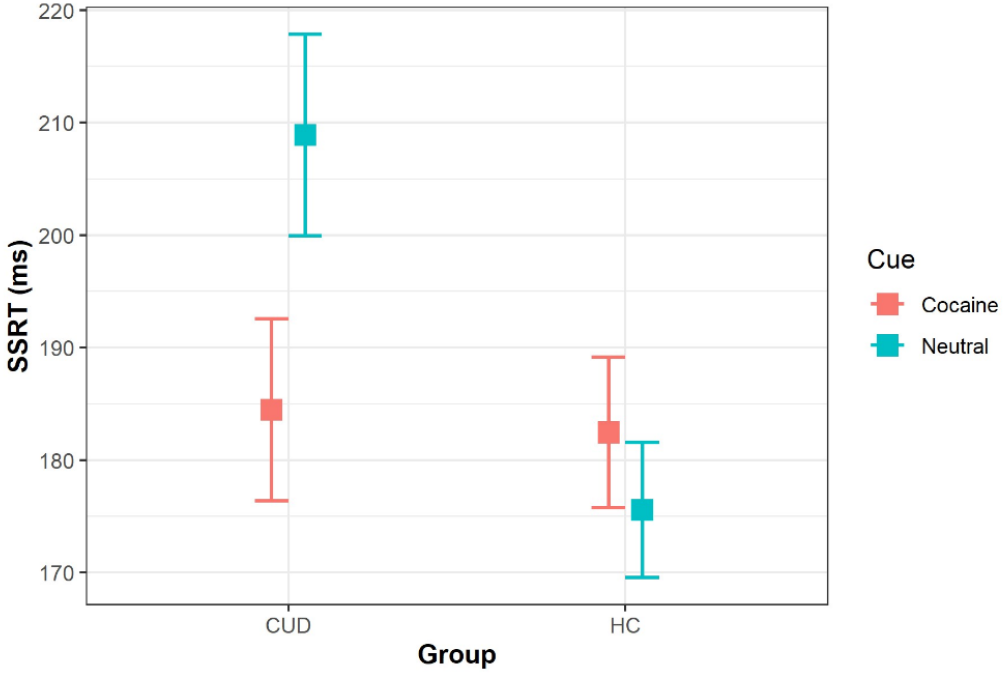
Stop Signal Reaction Times (SSRT). CUD participants were slower at stopping (higher SSRT) than HC when presented with neutral cues but were improved (faster SSRT compared to neutral cues (*p* = 0.035)) when presented with cocaine-related cues to reach the level of the HC participants, regardless of the observer conditions. No differences in SSRT between these types of cues were found in HC participants.

### 3.2 Neuroimaging results

The second-level analysis examining the neural signature of inhibitory control with GLM-Flex (contrast: Stop Correct > Stop Incorrect) led to significant Group X Social context X Cue type interactions in the bilateral OFC, right IFG, left MTG, and left visual cortex (V1) (*P*_*FWE*_ < 0.05). See Table 2.

**Table 2.** Whole brain results. Regions showing significant Group X Social context X Cue type interaction (Stop Correct > Stop Incorrect). (*P*_FWE_ < 0.05)

| Brain regions | Side | Cluster size | Peak MNI |  |  | Peak Z value | P-value cluster (corrected) |
| --- | --- | --- | --- | --- | --- | --- | --- |
|  |  |  | x | y | z |  |  |
| Orbitofrontal Cortex | Left | 166 | -40 | 54 | -10 | 4.55 | $p = 0.016$ |
| Orbitofrontal Cortex | Right | 151 | 40 | 56 | -14 | 4.36 | $p = 0.049$ |
| Inferior Frontal gyrus | Right | 133 | 56 | 27 | 28 | 4.43 | $p = 0.026$ |
| Middle Temporal gyrus | Left | 252 | -63 | -6 | -16 | 5.14 | $p = 0.001$ |
| Primary visual cortex | Left | 222 | -14 | -102 | -6 | 4.62 | $p = 0.003$ |

Subsequently, these significant interaction clusters were saved as separate masks using xjView (alivelearn.net/xjview/). Mean signal intensity from each cluster mask region was then extracted for each participant and task run. These values were entered into separate LMMs in R to explore which factor is driving the observed interactions. The same analysis was performed by focusing on the a priori defined ROIs in the bilateral STN. Results showed significant interactions in the right STN [χ^2^ (1) = 4.68, *p* = 0.030] (Figure 3), left OFC [χ^2^ (1) = 16.17, *p* <0.001], right OFC [χ^2^ (1) = 24.20, *p* < 0.001], right IFG [χ^2^ (1) = 19.28, *p* <0.001], left MTG [χ^2^ (1) = 18.19, *p* < 0.001], and left visual cortex (V1) [χ^2^ (1) = 20.02, p < 0.001]. See Figure 4. No significant results emerged in the left STN (*p* = 0.75).

#### CUD participants change in neural basis of inhibitory control as a function of the Social context and Cue type

Post-hoc analysis showed that during inhibitory control associated with cocaine cues (Cocaine > Neutral for Stop Correct > Stop Incorrect) there was a decreased activity in the right STN (Figure 2) when individuals with CUD were observed by the experimenter *versus* not observed (*Ps* < 0.001) (Figure 3). A similar effect was observed in left OFC, right OFC, and right IFG (Figure 4). Interestingly, an opposite pattern emerged in the left MTG in CUD participants, meaning an increased activity in this brain area in the observer condition compared to the non-observer condition (*p* < 0.001). The HC group showed no differences between these two conditions (all *Ps* > 0.08). Exact parameter estimates, t- and p values are provided in the Supplementary Materials (Tables S1-S5).

**Figure 3.**
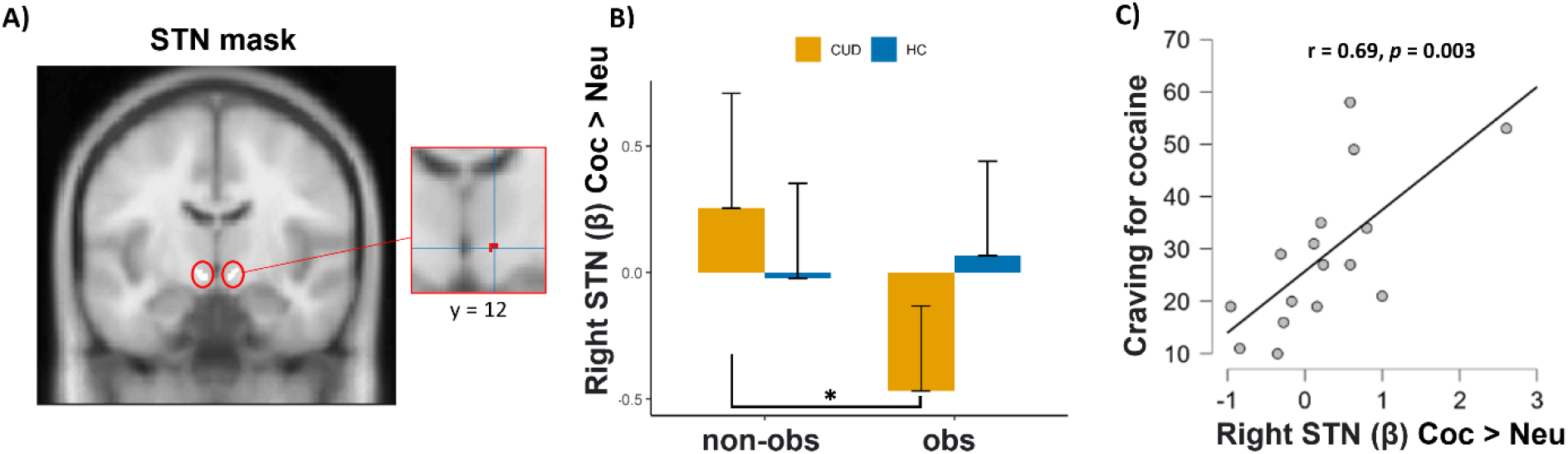
Brain Activations related to cocaine-related inhibitory control in presence or not of an observer. A) The left image is a bilateral STN mask defined by (60). The right image is a zoom-in inset showing significant Group X Social Context X Cue-type interaction within the right STN mask for the contrast Stop Correct > Stop Incorrect. B) bar graph showing reduced right STN activity during cocaine-related inhibitory control in CUD participants when being observed compared to when they were not (*p* < 0.001). No significant differences were found in the HC group (*p* = 0.775). Higher values indicate increased STN activity during inhibition for cocaine versus neutral cues. The error bars represent the standard mean of error. C) Craving for cocaine in CUD participants is positively associated with right STN activity during cocaine-related inhibitory control in the non-observer condition (r = 0.69, *p* = 0.003). **\***= *p* < 0.05.

**Figure 4.**
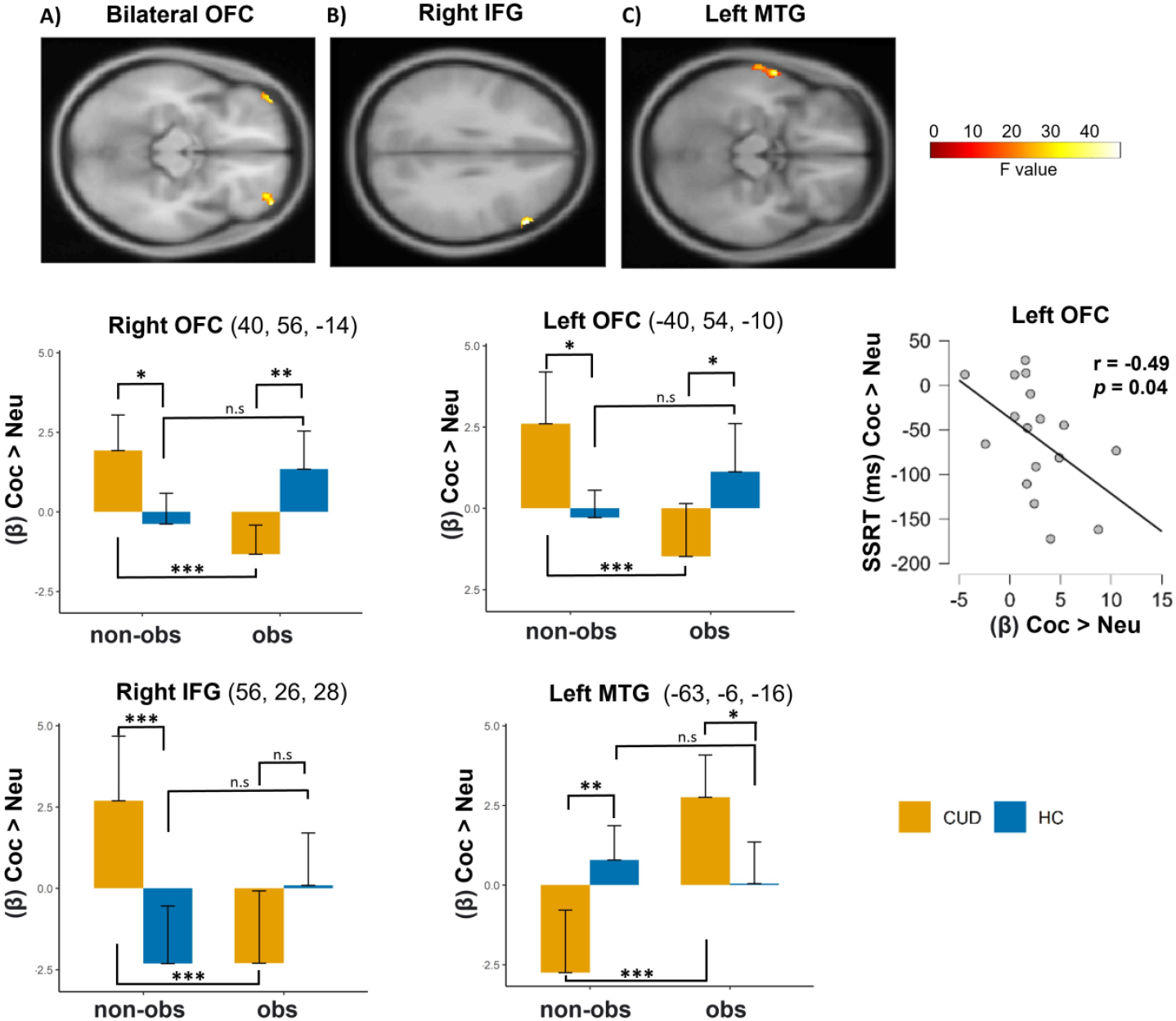
Brain activations related to cocaine-related inhibitory control in presence or not of an observer. Whole brain results showing significant Group X Social Context X Cue-type interactions in the left and right OFC (A), right IFG (B), and left MTG (C) (*P*_*FWE*_, < 0.05) for the contrast Stop Correct > Stop Incorrect. The bar graphs represent mean signal intensity from these significant interaction clusters. Higher values indicate increased activity during cocaine versus neutral inhibitory control. The scatterplot represents the significant negative correlation between CUD participants’ SSRT (cocaine > neutral) and left OFC activity for the same contrast (r = -0.49, *p* = 0.044). Lower values mean faster SSRT for cocaine compared to neutral-related cues. **\***= *p* < 0.05; **\*\***= *p* < 0.01; **\*\*\****p* < 0.001.

#### Differential effects of cue type and social context on inhibitory control: enhanced changes in CUD participants compared to HC

During cocaine-related inhibitory control (Stop Correct > Stop Incorrect), individuals with CUD showed higher brain activity in the left OFC, right OFC, and right IFG compared to HC when not observed (all *Ps* < 0.014). However, when observed by another person, CUD participants exhibited lower activity in the left OFC (*p* = 0.038) and right OFC (*p* < 0.001), indicating decreased brain activation during cocaine-related inhibitory control compared to HC. No significant differences were found in the right IFG between the CUD and HC groups in the observer condition (*p* = 0.156). Additionally, CUD individuals showed lower activity in the left MTG when not observed (*p* < 0.001) and higher activity when observed (*p* = 0.048) by another person compared to HC. Exact parameter estimates, *t*- and *p* values are provided in the Supplementary Materials (Tables S1-S5).

#### Craving for cocaine and SSRT are associated with STN and OFC activities during inhibitory control, respectively

Among participants with CUD, there was a significant positive correlation between craving for cocaine (CCQ scores) and activity in the right subthalamic nucleus (STN) during cocaine-related inhibitory control in the non-observer condition (r = 0.69, *p* = 0.003) (Figure 3C) but not in the observer one (*p* = 0.76). Moreover, in the non-observer condition, better inhibitory performance in CUD participants was significantly associated with higher activity in the left OFC (r = -0.49, *p* = 0.044) (Figure 4) and marginally associated in the right OFC (r = -0.48, *p* = 0.053). These correlations were not significant when participants were being observed (*Ps > 0*.*30)*. No significant correlations were found in the HC group. Lastly, linear regression analyses revealed that higher activity in the right OFC during cocaine-related inhibitory control (across observer/non-observer conditions) positively predicted increased activity in the right STN for both the CUD (β = 0.488, *p* = 0.003) and HC groups (β = 0.487, *p* = 0.003).

## 4. Discussion

By combining fMRI and a novel version of the stop-signal task, we investigated whether the control of inhibition in cocaine addiction can be affected by either cocaine-related cues and/or the social context and which are the neural correlates of this possible modulation.

Our study demonstrates that individuals with CUD exhibit a reduced ability to inhibit their responses in the SST, particularly when analyzing only trials associated with neutral cues to align with previous research that utilized traditional SST paradigms. This finding is in line with existing evidence indicating impaired inhibitory control among individuals with substance use disorders (37,39,40,42,59,61–63) including CUD (36,42,64). It underlines the significance of inhibitory control in addiction, as efforts to resist drug seeking and consumption rely heavily on voluntary, deliberate inhibition (34).

However, the engagement in seeking and taking drugs also depends upon the relative strength of the motivation or craving to use the drug (54,65), which can impact a person’s ability to control their impulses (41,66). To test the potential modulation of inhibitory control by drug cue salience, we included for the first-time cocaine-related cues (in forms of images) in an SST.

Interestingly, cocaine-related cues helped individuals with CUD to exhibit an improved inhibitory control compared to trials with neutral cues, reaching the level of HC. In contrast, HC did not show any difference in performance between cocaine and neutral-related trials, indicating that cocaine cues only affect inhibitory control in individuals with CUD and not in HC (possibly because drug cues were not salient or relevant for the HC). It might be surprisingly that cocaine cues did not impair the stopping performance of CUD participants, but rather improved it. One possible explanation for this finding is that these cues elicited craving for cocaine, potentially increasing motivation, arousal, and attentional focus on these salient trials specifically for CUD participants. Indeed, our neuroimaging results show that the more CUD participants experienced craving for reward, as measured by the CCQ questionnaire, the greater the activity in their right STN during cocaine-related inhibitory control.

Previous studies have shown the crucial role of the STN in inhibitory control processes, as a key component of the basal ganglia-thalamocortical circuit involved in motor and cognitive control (23–25,27). Also, the STN encodes both drug and natural reward values (16,33,67,68). Thus, our findings suggest that the presence of drug cue stimuli during an inhibitory task such as the SST can activate the STN, leading to increased inhibitory control, since STN inhibition reduces it (26,27,65).

Importantly, the STN receives direct inputs from cortical regions such as the IFG and the orbitofrontal cortex OFC via the hyperdirect pathway. It has been shown that these cortico-STN hyperdirect pathways facilitate the suppression of the ongoing motor response (23,28,29). Strikingly, our study revealed exactly the engagement of these brain regions during inhibitory control. In particular, CUD participants exhibited increased activity in these regions during inhibition in trials involving cocaine-cues compared to those with neutral cues. Conversely, HC participants showed heightened activity in these brain areas in trials involving neutral cues relative to cocaine cues. Our finding suggests that salient drug cues in addiction can modulate the activity of these frontal regions, which in turn may influence the STN, possibly via a network that regulates inhibitory control processes. This finding is further corroborated by the negative association that we found between the OFC activity during cocaine-related inhibitory control and CUD participants’ SSRT, showing that a higher activity in this brain area was associated with a better inhibition in trials with cocaine-cues. By highlighting the involvement of both OFC and STN in the regulation of the inhibition, our data confirm the network underscored in the review by (27).

As regards to the impact of the presence of an observer on inhibitory control, we found a reduced activity in the OFC, IFG, and STN in CUD participants under this condition. This decrease in activity could be attributed to the observer acting as an alternative reinforcer (9), dampening the salience and motivation associated with drug-related cues during the SST. In contrast, HC participants did not exhibit significant differences between observer and non-observer conditions, possibly because drug cues were not salient for them. Furthermore, it is worth noting that our study also revealed an interesting finding regarding the association between craving for cocaine, neural activity in the STN during cocaine-related inhibitory control, in line with the former findings showing that inhibition of the STN can reduce motivation for cocaine or escalation of cocaine intake in rats (16–18) and the relationship between OFC activity and stopping abilities in individuals with CUD. Importantly, these associations were significant only in the non-observer condition. This further suggests that the presence of an observer may dampen these associations, in line with the modulations observed after STN lesions in rats self-administering cocaine in presence of a peer (9). Showing that activity of STN is modulated by the presence of a peer further confirm the critical role of STN in the reduction of cocaine intake observed when subjects (rats like human) are in presence of a peer (9).

A further interesting neural signature of cocaine-related inhibitory control is the increased activity in the left MTG, when CUD participants were observed compared to when they were not. This difference was not found in HC participants. Also, when not under observation, individuals with CUD exhibited decreased activity in the left MTG as compared to HC. However, when observed by another person, their left MTG activity showed a significant increase compared to HC. The MTG is involved in various cognitive processes, including attention, perception, and social cognition (46,69–71). This heightened activity in the MTG could be a result of increased attention and vigilance triggered by the awareness of being observed. It may reflect anticipatory and evaluative processes where individuals focus on their actions and potential consequences in a social context.

Also, it is important to note that our study used the experimenter wearing a lab coat as the observer. Thus, the observer was a non-familiar person to the participant. Future human studies should explore the use of familiar peers on control of inhibition in CUD, as well as peers who are either drug-naïve or drug users, as this could yield different outcomes. Indeed, in a previous study on rats and human cocaine users(9), the drug consumption was reduced depending on the type of the peer, with a strong effect when a peer was present, abstinent, or drug-taking as well, further diminished when the peer was non-familiar.

To the best of our knowledge, this is the first study assessing the brain systems that regulate the complex interplay between drug cues, social factors, and inhibitory control in cocaine addiction in humans. Our findings can shed light on potential targets for intervention and suggest the importance of considering further the social context in addiction research and treatment. Given that there is still space for improvement in the management of cocaine-related disorders, these results may be crucial to develop harm reduction strategies for cocaine users.

## Financial support

This research was funded by the Centre National de la Recherche Scientifique (CNRS), Aix-Marseille Université (AMU), Institut de Recherche en Santé Publique (IRESP) and Alliance Aviesan in their call “IRESP-19-ADDICTIONS-02” and the support of the A*MIDEX project (ANR-11-IDEX-0001-02) funded by the “Investissements d’Avenir” French Government program, managed by the French National Research Agency (ANR).

## Competing interest

None.

## Notes

### Competing Interest Statement

The authors have declared no competing interest.

### Summary of Updates

A space was added between the Name and Surname of the first author

