## Supplementary materials for "Social Context and Drug Cues Modulate Inhibitory Control in Cocaine Addiction: involvement of the STN evidenced through Functional Magnetic Resonance Imaging" (from another version of this preprint: version 1, 2023-09-06)

| ***Comparison – left MTG*** | *Estimate* | *t-ratio* | *p-value* |
|---|---|---|---|
| Non-Observer Controls vs. Observer Controls | 0.679 | 0.659 | 0.9122 |
| Non-Observer Controls vs. Non-Observer Users | 3.502 | 3.432 | **0.0048** |
| Non-Observer Controls vs. Observer Users | -2.001 | -1.964 | 0.2089 |
| Observer Controls vs. Non-Observer Users | 2.824 | 2.771 | **0.0333** |
| Observer Controls vs. Observer Users | -2.680 | -2.626 | **0.0485** |
| Non-Observer Users vs. Observer Users | -5.503 | -5.416 | **<.0001** |

| ***Comparison – left OFC*** | *Estimate* | *t-ratio* | *p-value* |
|---|---|---|---|
| Non-Observer Controls vs. Observer Controls | -1.58 | -1.572 | 0.3990 |
| Non-Observer Controls vs. Non-Observer Users | -2.98 | -2.996 | **0.0179** |
| Non-Observer Controls vs. Observer Users | 1.12 | 1.131 | 0.6714 |
| Observer Controls vs. Non-Observer Users | -1.40 | -1.410 | 0.4962 |
| Observer Controls vs. Observer Users | 2.70 | 2.717 | **0.0384** |
| Non-Observer Users vs. Observer Users | 4.10 | 4.142 | **0.0004** |

| ***Comparison – right OFC*** | *Estimate* | *t-ratio* | *p-value* |
|---|---|---|---|
| Non-Observer Controls vs. Observer Controls | -1.766 | -2.431 | 0.0778 |
| Non-Observer Controls vs. Non-Observer Users | -2.331 | -3.084 | **0.0143** |
| Non-Observer Controls vs. Observer Users | 0.928 | 1.229 | 0.6101 |
| Observer Controls vs. Non-Observer Users | -0.566 | -0.749 | 0.8767 |
| Observer Controls vs. Observer Users | 2.694 | 3.563 | **0.0033** |
| Non-Observer Users vs. Observer Users | 3.259 | 4.552 | **0.0001** |

| ***Comparison –right IFG*** | *Estimate* | *t-ratio* | *p-value* |
|---|---|---|---|
| Non-Observer Controls vs. Observer Controls | -3.035 | -2.315 | 0.1015 |

| | | | |
|---|---|---|---|
| Non-Observer Controls vs. Non-Observer Users | -5.353 | -4.123 | **0.0005** |
| Non-Observer Controls vs. Observer Users | -0.292 | -0.225 | 0.9960 |
| Observer Controls vs. Non-Observer Users | -2.318 | -1.788 | 0.2852 |
| Observer Controls vs. Observer Users | 2.744 | 2.113 | 0.1561 |
| Non-Observer Users vs. Observer Users | 5.062 | 3.915 | **0.0009** |

| ***Comparison –right STN*** | *Estimate* | *t-ratio* | *p-value* |
|---|---|---|---|
| Non-Observer Controls vs. Observer Controls | -0.106 | -0.389 | 0.9798 |
| Non-Observer Controls vs. Non-Observer Users | -0.286 | -1.049 | 0.7211 |
| Non-Observer Controls vs. Observer Users | 0.437 | 1.604 | 0.3812 |
| Observer Controls vs. Non-Observer Users | -0.180 | -0.661 | 0.9114 |
| Observer Controls vs. Observer Users | 0.543 | 1.990 | 0.1988 |
| Non-Observer Users vs. Observer Users | 0.724 | 2.692 | **0.0409** |
